# Network-based analysis on pharmacological mechanisms underlying the anti-diabetic property of *Coix lacryma-jobi*

**DOI:** 10.1101/2021.10.15.464488

**Authors:** Arvin Taquiqui, Angelu Mae Ferrer, Janella Rayne David, Custer C. Deocaris, Malona V. Alinsug

**Author notes:** These authors contributed equally to this paper (Arvin Taquiqui, Angelu Mae Ferrer, Janella Rayne David).

## Abstract

Diabetes mellitus (DM) is the most common endocrine disorder and among the top 10 leading diseases causing death worldwide. Coicis semen [CS] (*Coix lacryma-jobi*), also known as adlay have been reported to display anti-diabetic properties. Unfortunately, studies on the pharmacological mechanisms involving adlay for the treatment of diabetes are nil. Thus, this study was conducted to evaluate the interactions and mechanisms of the bioactive compound targets of adlay in the treatment of diabetes using network analysis. Adlay bioactive compounds and potential target genes were obtained from SymMap. Diabetes related target genes were collected from CTD. Protein-Protein Interaction Network was analyzed using the STRING database. GO and KEGG pathway enrichment analyses were performed using DAVID to further explore the mechanisms of adlay in treating diabetes. PPI and compound-target-pathway were visualized using Cytoscape. A total of 25 bioactive compounds, 201 corresponding targets, and 35839 diabetes mellitus associated targets were obtained while 200 were considered potential therapeutic targets. The 9 bioactive compounds studied were berberine, oleic acid, beta-sitosterol, sitosterol, linoleic acid, berberrubine, jatrorrhizine, thalifendine, and stigmasterol. The identified 5 core targets were ESR1, JUN, MAPK14, and RXRA. Adlai targets enriched in GO terms were mostly involved with positive regulation of transcription, response to drugs, and negative regulation of apoptosis. This study provides novel research insights into the clinical properties of adlay in diabetes melitus treatment.

## Introduction

Diabetes mellitus (DM) is a chronic condition characterized by increased glucose levels with defects in secretion and action of insulin. Chronic hyperglycemia with disruptions of carbohydrate, protein, and fat metabolism results from having low levels of insulin. Diabetes is found to be linked with dysfunction and organ deterioration (American Diabetes Association, 2014). Moreover, type 2 diabetes mellitus (T2DM) is linked to a number of risk variables, including age, family history, calorie intake, hypertension, and obesity. Diabetes had a global incidence of 415 million in 2015, and it is predicted to hit 642 million by 2040. (Tripathy et al., 2017). However, current knowledge of the causes of diabetes and optimal treatment remains limited (Ge et al., 2018).

Natural products have long been used for the treatment of diabetes, particularly in Asia, Africa, and India. There have been numerous studies on herbal medicines for the discovery and development of antidiabetic medications. The hypoglycemic effects of variety of extracts and bioactive components have been studied. Natural product extracts are also commonly prescribed globally for diabetic therapy and many of these were investigated scientifically and to see what properties they have (Odhav *et al*., 2010). Coix (*Coix lachryma-jobi* L.), also known as adlay or Coicis Semen (CS), is an excellent source of traditional medicine and nutritious food that is primarily cultivated in China, Japan, and Burma. Coix is a grass that belongs to the Poaceae family and is a relatively close to maize. It is a Southeast Asian native that is cultivated for years in East and Southeast Asia (Zhu, 2017). The main medicinal component is the adlay seed, which contains a variety of bioactive substances including polysaccharides, polyphenols, coixol, lipids and proteins (Chiang et al., 2020). It has biological processes and functions such as anticancer capacity, antioxidation, and immune modulation based on recent epidemiological studies. It has a substantial antihypertensive impact and could be an excellent natural constituent of antihypertensive and related disorders (Li-Chun et.al, 2019). Since common knowledge about coix have received many attentions in discovering the scientific basis behind the claims of the health and medical effects it offers, this study assessed the potential role of coicis semen in the treatment of DM using network-based approach. Therefore, this study evaluated the interactions between the bioactive compound targets of *Coix lacyrma-jobi* L. and disease targets of DM and explored the mechanism of the bioactive compounds that may have therapeutic mechanism for DM.

## Materials and Methods

### 2.1 Information acquisition of compound composition of Coicis semen (CS) and Diabetes Mellitus (DM)

The compound composition of Coicis semen seed were retrieved from the ‘ingredient tab’ in SymMap (http://www.symmap.org/, Version 2), a database that provides massive information on herbs, ingredients, targets, as well as clinical symptoms and diseases (Wu et al., 2019). Data acquisition of adlay was done using ‘*Coicis semen’* as the keyword. Other information of the obtained components including, molecular weight, PubChem ID, and canonical smiles were confirmed and obtained using PubChem database (https://pubchem.ncbi.nlm.nih.gov/). To construct a network with Diabetes Mellitus, target genes of the disease with the same name as the keyword were searched and obtained from the Comparative Toxicogenomics Database (CTD, https://ctdbase.org/, updated 2^nd^ September 2021) through the ‘gene’ tab and with limitations to disease identifier. CTD is a manually curated information about the integrated interactions of chemicals, genes, phenotypes, diseases, and how environmental exposures affects human health (Davis et al., 2020). Information obtained from these databases were imported and organized in separate sheets to avoid mix-ups of data and were saved as TSV files.

### 2.2 Acquisition of Bioactive Compounds through Pharmacokinetic ADME Evaluation

All retrieved compounds were screened with oral bioavailability (OB) value and drug-likeness (DL). Initially, compounds were retained only if OB ≥ 30 (Wang et al., 2017). Compounds without OB information were also removed from the list. Then, using the Swiss ADME website (http://www.swissadme.ch/) the retained compounds were screened with DL with limitation to the Lipinski’s Rule of Five (RO5) by inputting the Canonical SMILES corresponding to each compound. A compound with more than one violation were removed from the list. Compounds that satisfied the criteria were identified as potential bioactive compounds.

### 2.3 Intersection of bioactive compound potential target and disease related target genes

The obtained Coicis Semen-related target genes and Diabetes Mellitus-associated target genes were separately imported into Microsoft (MS) excel spreadsheet to remove duplicate genes. After that, both of the target genes were combined into a single column and duplicates were screened and copied into a different sheet. From that, duplicates were then removed to obtain the common target genes.

### 2.4 Protein-Protein Interaction (PPI) Network Analysis

The common target genes of Coicis Semen and Diabetes Mellitus were uploaded to Search Tool for the Retrieval of Interacting Genes (STRING, https://string-db.org/, Version 11.5) as the background network database with limitation to “*Homo sapiens”* and a highest confidence score of 0.900 to construct a high-quality and credible PPI network (Zhou et al., 2021). The target protein interaction was obtained and saved as a TSV file. Then the TSV file was imported into Cytoscape software (http://www.cytoscape.org/, ver 3.8.2) to analyze and visualize the PPI network. Using the Network Analyzer tool, summary statistics of the nodes and edges, along with the topological parameters such betweenness centrality, closeness centrality, and the degree values of each target were obtained.

### 2.5 Enrichment Analysis and Network Construction

The Database for Annotation, Visualization, and Integrated Discovery (DAVID, https://david.ncifcrf.gov/home.jsp, version 6.8) was used for GO and KEGG pathway functional enrichment analysis. By starting the analysis, the Gene symbol of common targets of “CS-DM” was imported in the gene list. OFFICIAL_GENE_SYMBOL, *Homo sapiens*, and gene list were selected in the identifier, species, and list type, respectively. The screening criteria was limited to P ≤0.05 using Bonferonni correction (Liang et al., 2014). Biological process (BP), Cellular Component (CC), and Molecular Function (MF) are the three components of GO term enrichment analysis. GOTERM_BP_DIRECT, GOTERM_CC_DIRECT, and GOTERM_MF_DIRECT, as well as KEGG_PATHWAY were selected in Gene Ontology and Pathways in the annotation category, respectively to analyze and visualize the results. Pathways with the top 20 protein numbers were used for the establishment of the compound-target-pathway network by Cytoscape. A summary of the workflow is presented in Figure 1.

**Figure 1.**
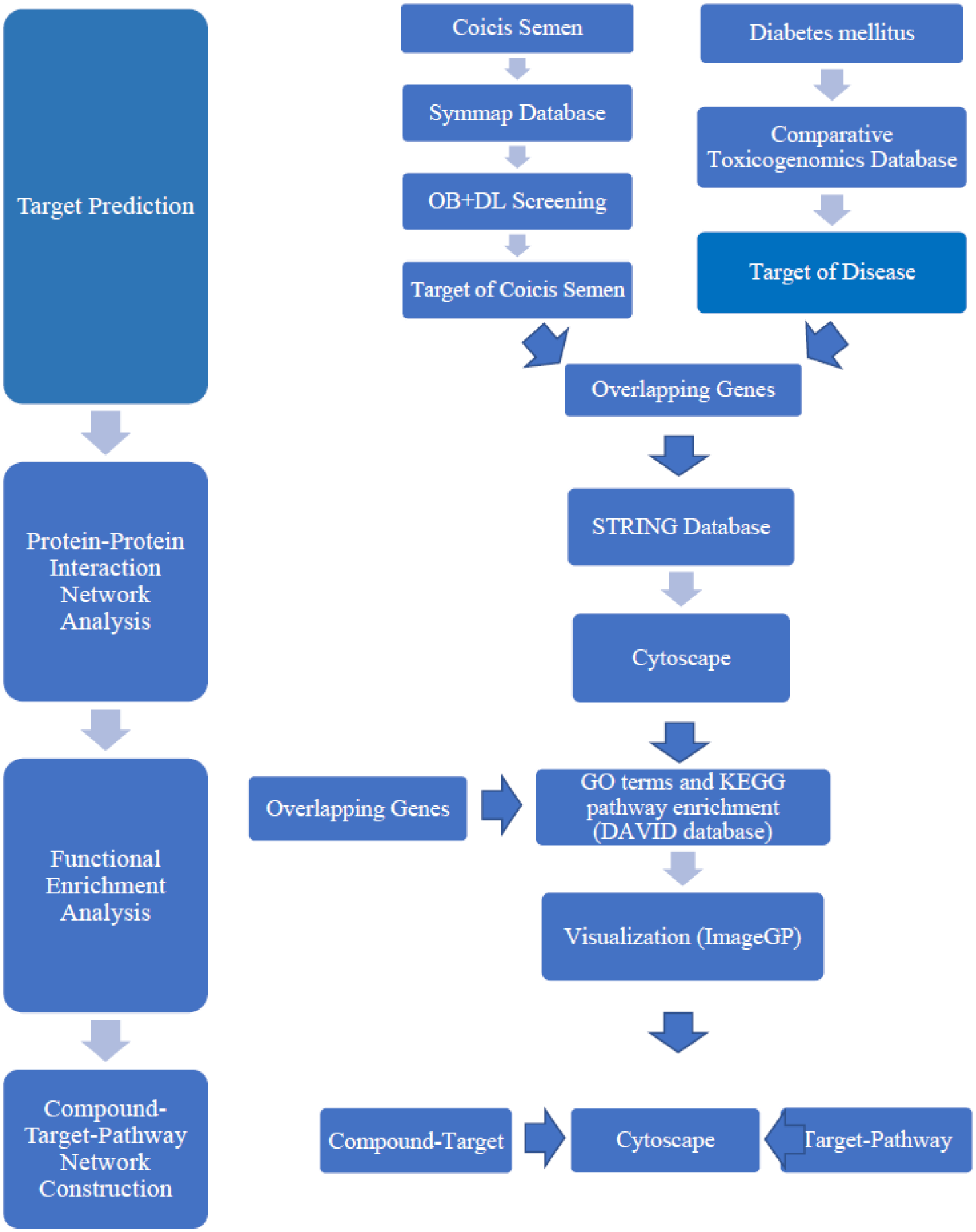
Schematic workflow. Summary of methods conducted for the network pathway analysis of *Coix lachrymal jobi* components against diabetes is presented.

## Results

### 3.1 Compounds, Corresponding Targets, and Genes of Coicis Semen

There were 111 compounds of Coicis Semen obtained from SymMap database. 226,127 target genes were related to Diabetes Mellitus from CTD, and a total of 507 were identified as potential target genes of CS from SymMap.

### 3.2 Collection of Bioactive Compounds and Targets of Coicis Semen using ADME Evaluation

Frequently, drugs are directed orally in clinical medication (Li et al., 2021). Thus, it is essential that from the selection up to identification to be an effective treatment for the patient, a compound must exhibit high biological activity and low toxicity (Daina et al., 2017). Daina et al. (2017) claimed that the initial assessment of ADME (Absorption, Distribution, Metabolism, and Excretion) in the discovery phase exhibited a significant reduction of the failures associated with pharmacokinetics in the clinical phases. In keeping with the Traditional ADME screening principle (Xu et al., 2020), Oral Bioavailability (OB) and Drug Likeness (DL) are the two main variables used for the screening of compounds that affect drug absorption across the gastrointestinal mucosa (Huang et al., 2020).

OB is the intensity and rate of absorbing drugs into the systemic circulation. The value of the percentage of the OB of the compound is directly proportional to the probability of the clinical application development of the compound (Chen et al., 2018). High OB is normally a core indicator in the establishment of the DL index of bioactive molecules (Huang et al., 2020). Drug-likeness represents the capacity of molecules in the compound demonstrating physicochemical properties comparable to existing drugs (Potunuru et al., 2019). Lipinski’s Rule of Five (RO5) is an empirical guideline that predicts the high risk of poor absorption or permeability of a compound on its underlying simple physicochemical properties. For a compound to be considered an orally available drug, it must satisfy the following conditions; (1) no more than 5 hydrogen-bond donors, (2) no more than 10 hydrogen-bond acceptors, (3) the molecular weight (MWT) is less than 500 Dalton, and (4) the calculated Log P (CLogP) or lipophilicity is less than 5 (A decade of Drug likeness, 2017). A compound with 2 or more violations of these conditions is considered to be a non-orally available drug (Lipinski et al., 2001).

The screening condition for OB was set as ≥ 30 (Chen et al., 2018). For DL, a compound must have no more than 1 violation from the Swiss ADME. A compound that satisfied both criteria was considered a bioactive compound. From the 111 compounds retrieved from SymMap, 30 were with oral bioavailability greater than or equal to 30. 2 of the compounds were unable to be found in PubChem and have no PubChem ID while 1 have no target genes. Finally, after ADME evaluation and potential target gene prediction, a total of 25 compounds were identified as potential bioactive. Detailed information on Coicis semen compounds is presented in Supplementary Tables S1.

The corresponding target genes of each identified bioactive compound were obtained from SymMap. From the 507 target genes, a total of 201 target genes were obtained after the removal of duplicates. Detailed information on Coicis semen-related targets is presented in Supplementary Table S2.

### 3.3 Identification of Targets Related to Diabetes Mellitus

CTD was mainly used to identify target genes related to Diabetes Mellitus. From 226,127 potential related disease targets genes, a total of 35,839 target genes associated with DM were identified after the removal of duplicate targets. Detailed information on DM-related targets is presented in Table S4.

The bioactive compound targets and disease targets were intersected to obtain 200 common targets as shown in Figure 2. These common targets were taken as the central target for subsequent enrichment analysis and protein-protein interaction network.

**Figure 2.**
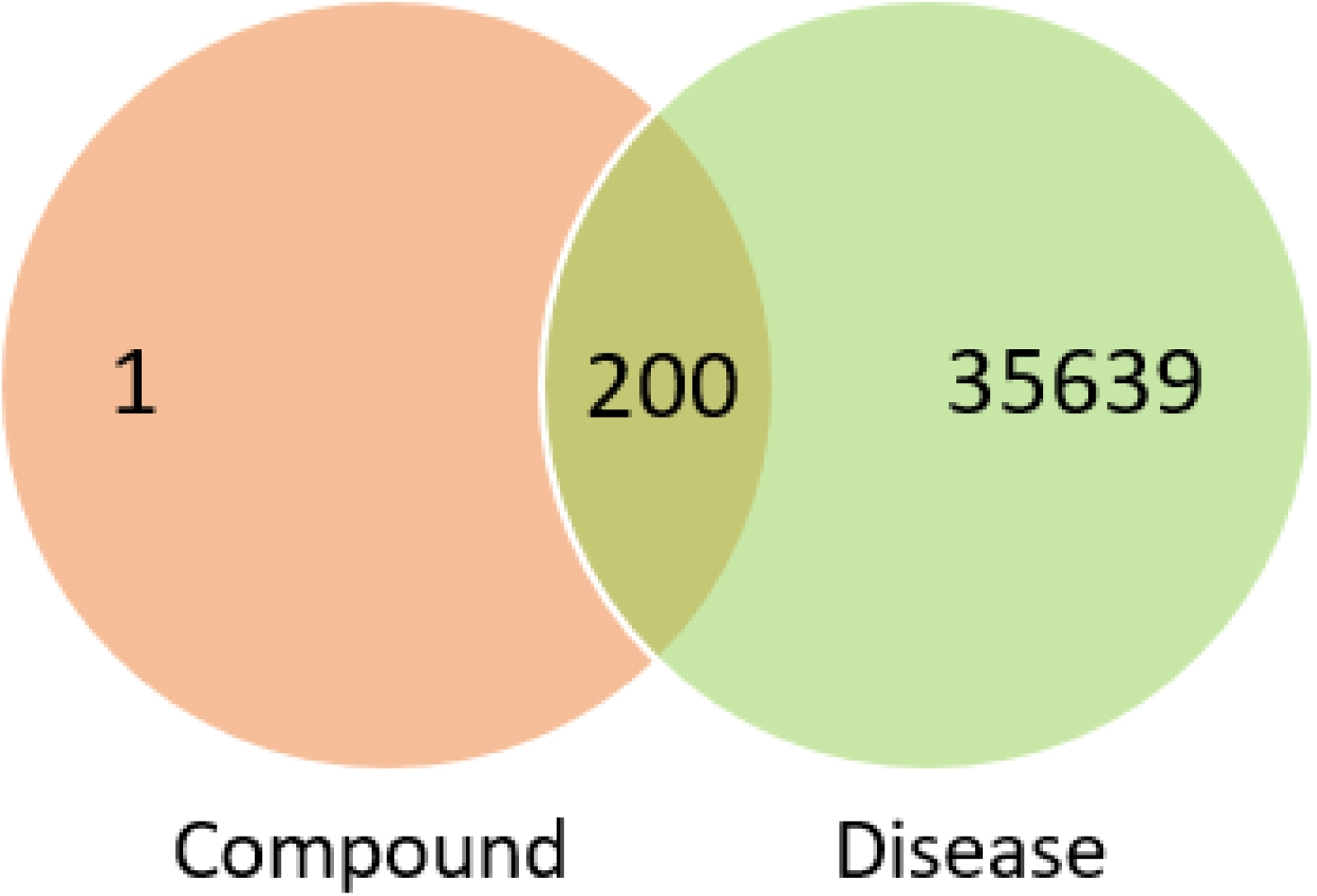
Venn diagram of target of coicis semen compound and diabetes mellitus disease

### 3.4. Protein-Protein Interaction Network

The PPI Analysis of 200 common targets of Coicis semen and Diabetes was executed through STRING database with limitation to “*Homo Sapiens*” and with a confidence score of 0.900. The PPI network then was established and visualized using Cytoskape 3.8.2, of which results (Figures 3-4) generated a total of 153 nodes representing all the predicted targets; with 652 edges representing the correlation between the active components of Coicis Semen and potential Diabetes targets. Network Analyzer tool in Cytoskape was utilized for the calculation of the topological parameters of the PPI network to identify hub nodes and essential targets wherein nodes with higher betweenness centrality were depicted by darker color in the network. Based on the result from the network analyzer, the top 10 targets of Coicis-Diabetes; MAPK3, JUN, MAPK1, TP53, RXRA, RELA, NCOA1, MAPK14, ESR1 and FOS as defined by the highest degree and were described by a larger size in the network. These were then identified as hub nodes and essential targets of the PPI network (Supplementary Table 5).

**Figure 3.**
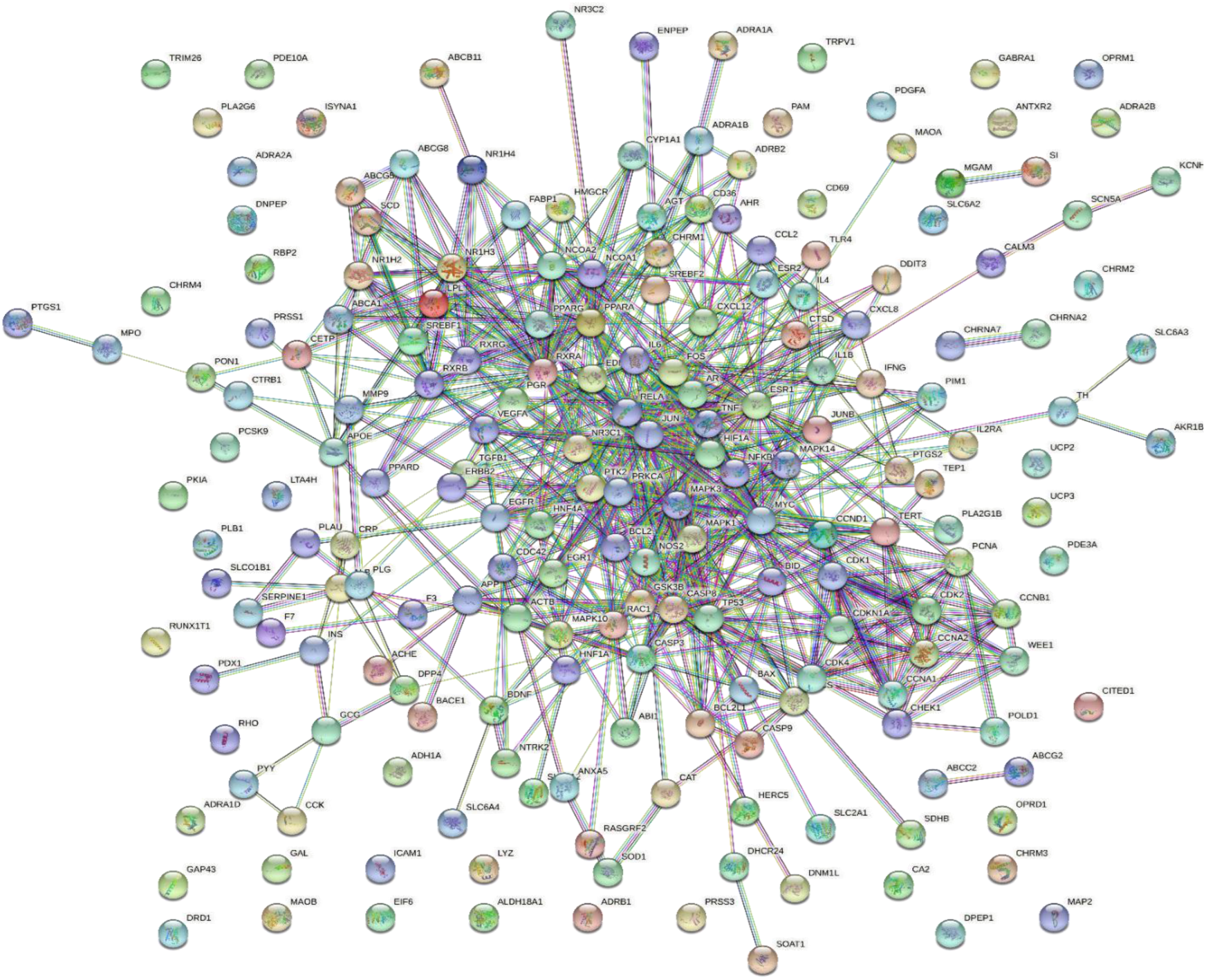
Interaction relationship between target proteins of coicis semen in diabetes was inferred using String. It is noteworthy that the common target protein of coicis semen is liver cancer. PPI enrichment is set at a p value <1.0e16.

**Figure 4.**
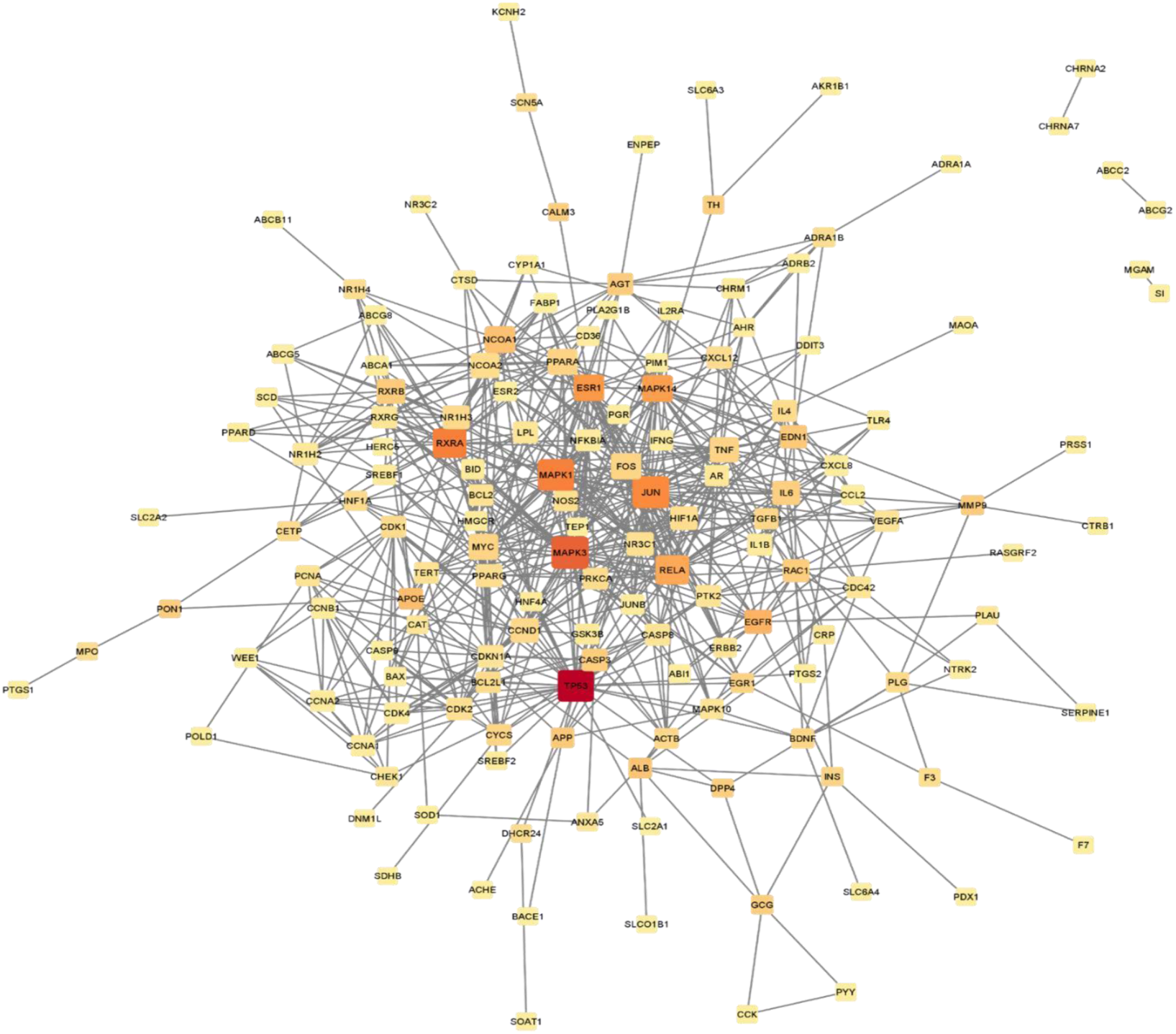
Interaction network of potential proteins of coicis semen in treating diabetes was inferred using Cytoscape composition results.

### 3.5. GO and KEGG Pathway Enrichment Analysis

The GO and KEGG Pathway Enrichment Analysis using the 200 common targets of coicis semen and Diabetes mellitus was executed through the DAVID database to identify potential mechanisms of Coicis semen against diabetes with a p-value cut-off of 0.05 as the screening condition. Bonferroni values were used for the identification of the top 20 significantly enriched GO terms and KEGG pathways of the gene overlaps (Figures 5). Biological Process (BP), Cell Component (CC) and Molecular Function (MF) were used as the main components of the GO term enrichment analysis and KEGG for the pathway enrichment analysis. According to the results, the 200 targets were significantly enriched in 500 BPs, 135 CCs, 109 MFs and 114 pathways. In the biological process (BP) category, the target proteins were mainly involved in positive regulation of transcription from RNA polymerase II promoter, response to drug, positive regulation of transcription, DNA-templated, and negative regulation of apoptotic process. In the cellular component (CC) category, the target proteins were mainly involved in cytoplasm, plasma membrane, cytosol, and extracellular space. Molecular Function (MF) analysis has a higher enrichment of protein binding, transcription factor activity, and sequence-specific DNA binding. The KEGG enrichment analysis results showed that pathways in cancer was the most closely related to the function of CS in Diabetes Mellitus treatment.

**Figure 5.**
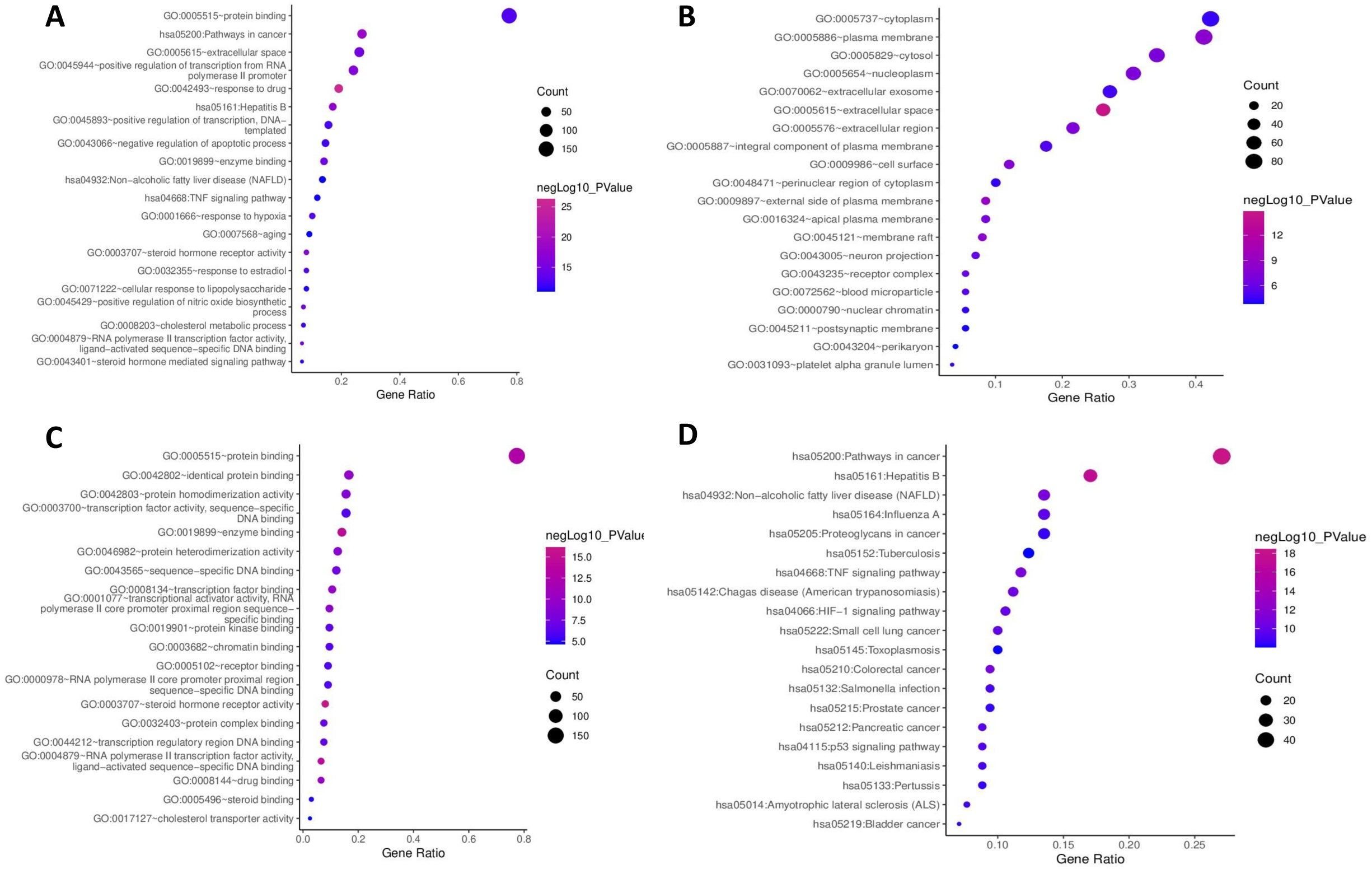
Gene Ontology and KEGG pathway analysis. Enrichment results were screened from 4 main components, namely: (A) Biological Processes (BP); (b) Cellular Components (CC); (B) Cellular component (C) Molecular Functions (MF); (D) KEGG pathways.

**Figure 6.**
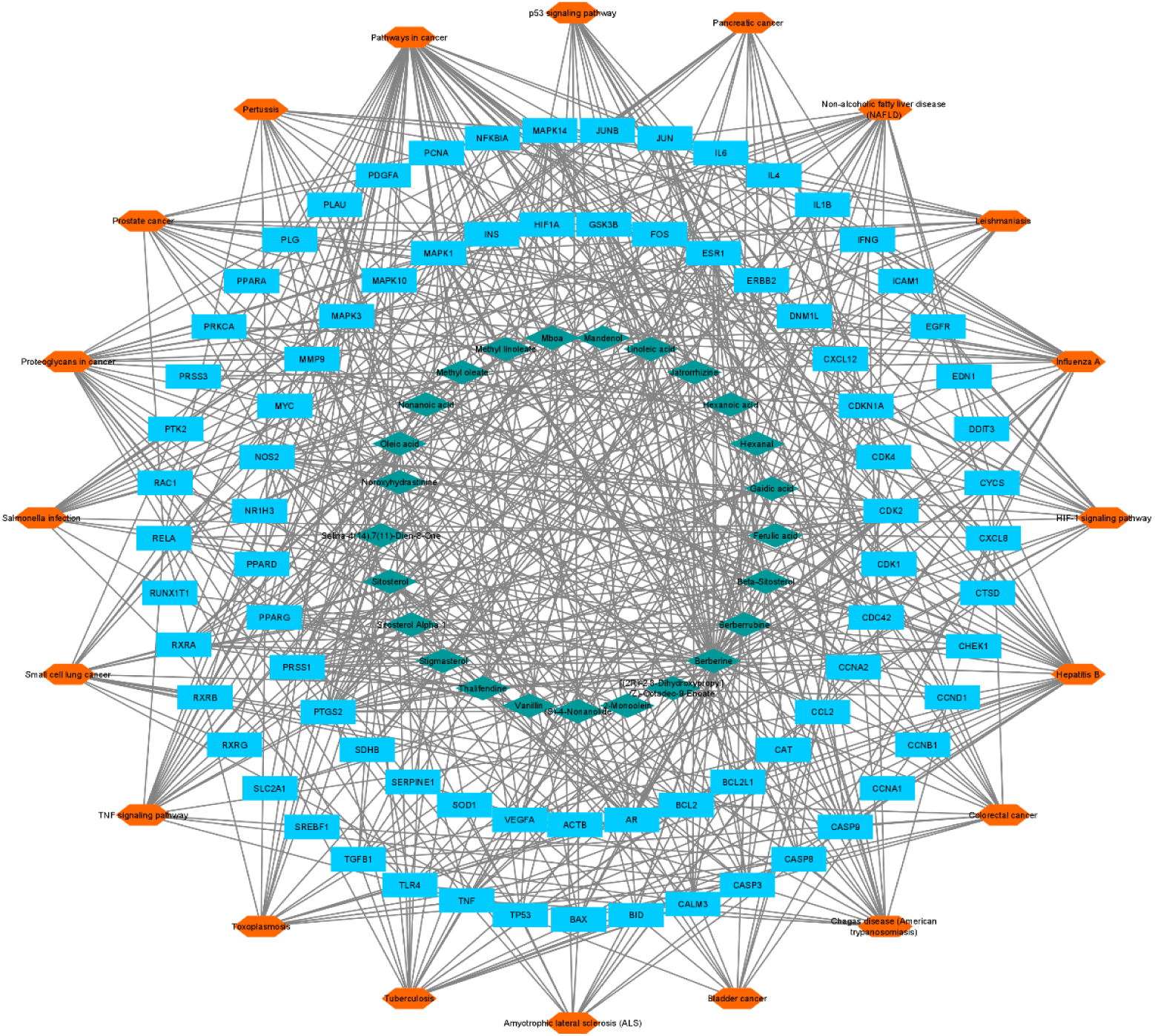
Compound-Target-Target Pathway Network. Green diamonds represent bioactive compounds of Coicis Semen, blue rectangles represent target genes, and orange hexagons represent the top 20 pathway.

### 3.6. Construction of Compound-Target-Target Pathway Network

The constructed compound-target-pathway network was established using Cytoscape 3.8.2 according to KEGG pathway enrichment result. The compound-target-pathway included 125 nodes and 607 edges, green diamonds represent bioactive compounds of Coicis Semen, blue rectangles represent target genes, and orange hexagons represent the top 20 pathway. Edges represent the relationship between compound and target, and between target and pathways. Hence, targets are bridges between compounds and pathways. In the network, 9 compounds had a higher-than-average degree, which revealed that they performed a key role in the compound-target-pathway network.

The compounds were berberine, oleic acid, beta-sitosterol, sitosterol, linoleic acid, berberrubine, jatrorrhizine, thalifendine, stigmasterol. The top 8 hub nodes and essential targets of the PPI network were identified based on the overlap of the top 20 targets of degree, between centrality, and closeness centrality, which were ESR1, JUN, MAPK1, MAPK14, MAPK3, RELA, RXRA, and TP53. The interaction of the top 8 targets of the PPI network and the compound-target-pathway network was identified as core targets, indicating that they play a significant role in both PPI network and compound-target-pathway network. Lastly, four targets, ESR1, JUN, MAPK14, and RXRA were identified as core targets.

## Discussion

Following heart disease, cancer and other chronic illnesses, diabetes mellitus (DM) is among the top 10 leading diseases causing death worldwide. As characterized by hyperglycemia, it is a secretory and metabolic disorder known to affect carbohydrates, lipids and protein metabolism causing various complications in the body resulting to long-term dysfunction and organ failures mostly involving the eyes, heart and blood vessels, kidneys and even nerves. DM is also ascribed to defects in insulin secretion, insulin action or both (ADA, 2005), along with other factors. Bataski (2005) reported that diabetes mellitus is considered as the most common endocrine disorder in which more than 300 million people will be diagnosed and suffering with the disease by the year 2025. This chronic metabolic disorder is a fast-growing worldwide problem having a massive consequence in social, health, and economic aspects (Kaul et al., 2013). Moreover, diabetes related complications, if not treated, can be lethal (Suji & Sivakami, 2003).

Due to this, various treatments including insulin sensitizers, insulin-secreting drugs and insulin are being administered to combat the multifactorial effects of DM, however it is of without numerous side effects. (Li et al., 2019). Consequently, efforts in finding alternative treatments with comparable therapeutic efficacy is being mostly considered. This study explored the regulatory mechanism of the constituents of Coicis semen in the treatment of Diabetes Mellitus through network pharmacology. Based on various screenings, a total of 201 bioactive compounds of Coicis semen were obtained for this analysis from the SymMap database. Subsequently, this study obtained 200 target genes by acquiring the gene overlaps of the bioactive compounds of Coicis semen and the DM-associated genes acquired from the CTD database.

According to the PPI network analysis, constituents of Coicis semen are closely related to various target proteins in DM inferring that the therapeutic efficacy of Coicis semen in the treatment of DM relies on the combined action of several target proteins rather the regulation of a single target protein. With this, the top 10 targets of Coicis-diabetes with the highest degree based on the PPI network were selected which include MAPK3, JUN, MAPK1, TP53, RXRA, RELA, NCOA1, MAPK14, ESR1, and FOS1.

Moreover, the enrichment results revealed that 500 BPs, 135 CCs, 109 MFs and 114 pathways were involved. 9 bioactive compounds and 5 core targets were identified based on the topological parameters of the PPI network and compound-target-pathway network. The 9 bioactive compounds were Berberine, Oleic acid, Beta-Sitosterol, Sitosterol, Linoleic acid, Berberrubine, Jatrorrhizine, Thalifendine, Stigmasterol. The 5 core targets were identified as ESR1, JUN, MAPK14, and RXRA which could play a crucial role in the pathway of treating DM. Pathways in Cancer and Non-alcoholic fatty liver disease were among the pathways being targeted by these genes, which are most closely related to the function of Coicis Semen in treating Diabetes. Meaning, Coicis Semen may exert anti-diabetic by these targets.

MAPK1, MAPK3 and MAPK14 of the Mitogen-activated protein kinase (MAPK) family involve mechanisms that regulate a number of biological processes in response to various cellular signals such as insulin-signaling. The oxidative stress-activated p-38a, encoded by the MAPK14 gene is thought to be involved with decreased insulin-dependent stimulation of insulin signaling components and glucose transport activity which might induce potential role in the gene module response to hyperglycemia (Tang et al., 2012). Additionally, p38a has been shown to act in the promotion of energy expenditure, regulation of glucose homeostasis and lipid metabolism and management in physiological and pathophysiological states as demonstrated in several studies (Sumara G et al., 2009; Barger PM et al., 2001).

RXRA is a retinoid-X receptor that is involved in various metabolic processes such as glucose regulation and lipid metabolism (Pan et al., 2014). Several studies also revealed that its processes are closely related to diabetes (Franzago et al., 2019; Castello-Castillo et al., 2019). Firstly, as treatments for DM including novel therapies improving insulin action involves ligands that bind this retinoid X receptor wherein it functions by regulating transcription of genes that are associated in insulin action, adipocyte differentiation and lipid metabolism through heterodimers. Moreover, selective ligands of RXRA play significant functions by improving metabolic defects associated with DM especially T2DM such as hyperglycemia hyperlipidemia, and insulin resistance implying RXRA as a molecular target for Diabetes. (Lenhard, 2001).

As known, ESR1 has been demonstrated as a modulator of the glycemic homeostasis and is associated with metabolic disorders such as obesity, insulin-resistance, blood pressure; and type-2 diabetes (Huang et al., 2006; Barreto, JN. et al., 2018; Gallagher et al., n.d). Study of Qui et al. (2018) further demonstrated that ESR1 provides a functional role in glucose and lipid metabolism suggesting that ESR1 signaling pathway could be a promising therapeutic target for the prevention of the lipid and glucose metabolic defects.

Beta-Sitosterol is one of the several and principal phytosterols in plants with a chemical structure similar to that of cholesterol (Rashed 2020; Van, 2000). Beta-Sitosterol has been reported to have antidiabetic activity by increasing the fasting plasma insulin levels, decreasing fasting glycemia, and reducing glucose levels (Saeidnia, 2014) and anticancer activity including breast cancer, prostate cancer, colon cancer, lung cancer, stomach cancer, and ovarian cancer (Rashed, 2020). Studies suggested that Beta-Sitosterol is an effective apoptosis-promoting agent and that integration in the diet could provide a preventive measure for breast cancer (Awad et al., 2007).

Aside from anticancer activities, Berberine is also known to have anti-diabetic properties as it has been reported to function in the regulation of glucose and lipid metabolism. Additionally, several studies reported that Berberine could help type-2 diabetes patients by regulating blood sugar. It was also shown that Berberine can function in lowering blood insulin levels among newly-diagnosed type2 diabetic patients by enhancing insulin sensitivity. Aside from this, Berberine shows properties in insulin resistance reduction which is the indicator of type 2 diabetes. Several studies also conclude that Berberine could be a comparable alternative treatment to metformin which is commonly used to treat type 2 diabetes and offer better result in reducing blood glucose. Given the high potential of Berberine as an emerging antidiabetic agent, backed by animal and human trials, large-scale trials must still be acquired for further evaluation of the therapeutic efficacy of Berberine on Diabetes specially in the T2DM subtype.

Diabetes Mellitus along with other complex chronic diseases are a huge danger to the well-being and existence of humans. Thus, considered as one of the extremely critical challenges the world is facing (Shi et al., 2014). However, a significant reduction in the degree that fresh drug candidates are being transformed into clinical treatments with a high efficacy rate and value has been evident over the last few decades (Hopkins, 2008). With the breakthrough and rapid advancement of fields of bioinformatics, systems biology, and polypharmacology (Li & Zhang, 2013; Shi et al., 2014; Zhang et al., 2019), the concept of network pharmacology was presented by Andrew L. Hopkins (Shi et al., 2014) in order to examine the effect and intermediation of drugs as well as to demonstrate the synergistic principle of multicomponent drugs to search for multiple target treatment having high efficacy and low toxicity activity rather than single target (Tang et al., 2016; Zhang et al., 2013). Moreover, network pharmacology, by means of a systematic approach, examines on the interactions between drugs, targets, diseases, and compound-proteins/genes-disease and visually displays the network of drug targets. Still, it explored the influence of drugs on the biological network from a holistic perspective and is able to explain complexities among biological systems, drugs, and diseases from a network perspective (Zeng & Yang, 2017; Lui &Bai, 2020; Zhang et al., 2019). Consequently, network pharmacology is of great help in identifying the mechanism of action and assess the efficacy of drug through discovering the influence of compounds to the biological network (Gu et al., 2013).

Network pharmacology highlights the change of paradigm from the traditional “one target, one drug” to “network target, multicomponent therapeutics,” drug discovery (Li et al., 2014). According to Zhang et al. (2013), the advantages of network pharmacology study involve: (1) regulation of the signaling pathway with multiple channels; (2) increase in the efficacy of drug; (3) decrease of side effects; (4) increase in the success rate of clinical trials and; (5) reduction in the financial costs in drug discovery. As a result, this network-based drug discovery is deemed a promising method and has great potential to become the next generation approach of drug discovery (Li et al., 2014).

With the rapid advancement of research, network pharmacology, including the investigation of traditional medicines has become a new and prevalent method for drug mechanism research and drug development (Zhang et al., 2019; Lee et al., 2018). Numerous efforts have been made to employ the method to examine the complex of ingredients, unknown targets, as well as pharmacological mechanisms of herbal formulas to elucidate and provide understanding into the complex mechanisms of herbal formulas used to treat complicated diseases (Lee et al., 2018).

Through scientific exploration, the elucidation of various biological processes of adlay has been established despite the existence of its therapeutic application for thousands of years. Hence, it paved way for the occurrence of integration of traditional and modern medicine (Kuo, 2012) which was also evident from the different bioinformatics tools. Traditionally, adlay seed has been utilized as herbal medicine for the treatment of diabetes, asthma, inflammation, cough, diarrhea (Choi et al., 2015), pulmonary edema, wet pleurisy, chronic ulcer, dysuria (Feng et al., 2020), rheumatism, osteoporosis, (Zhang et al., 2019) chronic gastrointestinal disease (Choi et al., 2015; Feng et al., 2020), and as a diuretic (Zhang et al., 2019, Zhu, 2017; Wu et al., 2014).

A study on the biological activities of proteins from adlay had shown positive effects in the improvement of type II diabetes in mice (Watanabe et al., 2012). An examination conducted by Chen et al. (2019) concluded that the management of polysaccharides from adlay seed in diabetic mice exhibited an improvement in polydipsia, and showed a reduction in the loss of body weight, blood glucose, and insulin levels. Similarly, recent examination on the biological functions in mice revealed that the adlay seed polysaccharide possessed anti-diabetic properties as it decreased blood glucose levels, enhances glycemic control with no reduction in body weight, and behaved as an insulin secretagogue. Hence, could be exploited for anti-diabetic drug (Chen et al., 2019). However, despite the evidence provided by these reports, there have been insufficient studies that investigate the pharmacological mechanism of coicis semen in the treatment of diabetes mellitus which would be an essential aspect in the development of a drug. Consequently, the present study could present elucidation, reference, and guidance in the defects and later clinical research and development (Shen et al., 2021).

## Conclusion

Overall, this study assessed the therapeutic potential of Coicis semen in treating Diabetes Mellitus through a network-based analysis based on the aspect of constituents, targets and pathways. 201 compounds in Coicis semen were identified as potentially bioactive compounds that could target numerous DM-associated genes and facilitate therapeutic property. Based on the results of enrichment analysis, Coicis semen targets were enriched in GO terms mostly involved with positive regulation of transcription, response to drug, negative regulation of apoptotic process. Herewith, our findings offer insights into the clinical properties of Coicis semen in DM treatment and suggest that further clinical trials, pharmacological methods and research must be established to verify findings of this study for better understanding of the antidiabetic potential of adlay that could pave a way for drug development.

## Acknowledgements

The authors are grateful to the unwavering support of the Biology Department faculty of Polytechnic University of the Philippines. Special mention goes to Prof. Lourdes Alvarez, Prof. Arci Bautista, Prof. Julie Charmain Bonifacio, and Prof. Chester C. Deocaris for their guidance, patience, and moral support.

## Author Contribution

MVA and CCD conceptualized & designed the study, interpreted the data, and edited the manuscript; AT, AMF, and JRD performed data mining, analysis, and wrote the draft manuscript. All authors read and approved the final manuscript.

## Data Availability

The data supporting the findings of this study are available in the paper and from the corresponding author, Malona V. Alinsug, upon request.

## Ethics Approval

NA

## Consent to Publish

All authors read and approved the final manuscript.

## List of Tables

**Table 1.**
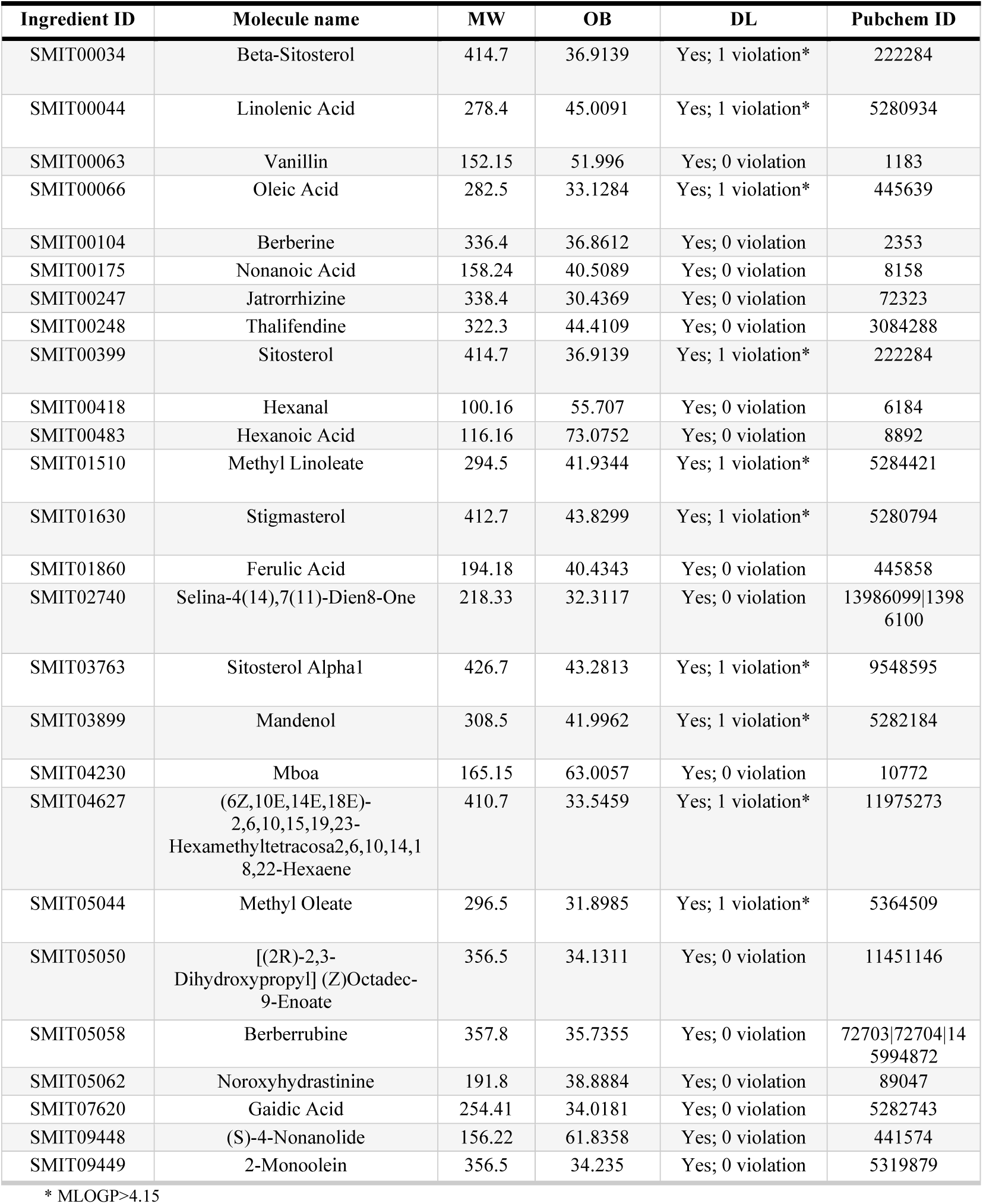
**L**ist of selected active compounds from coicis semen for network analysis

**Supplementary Table S1.**
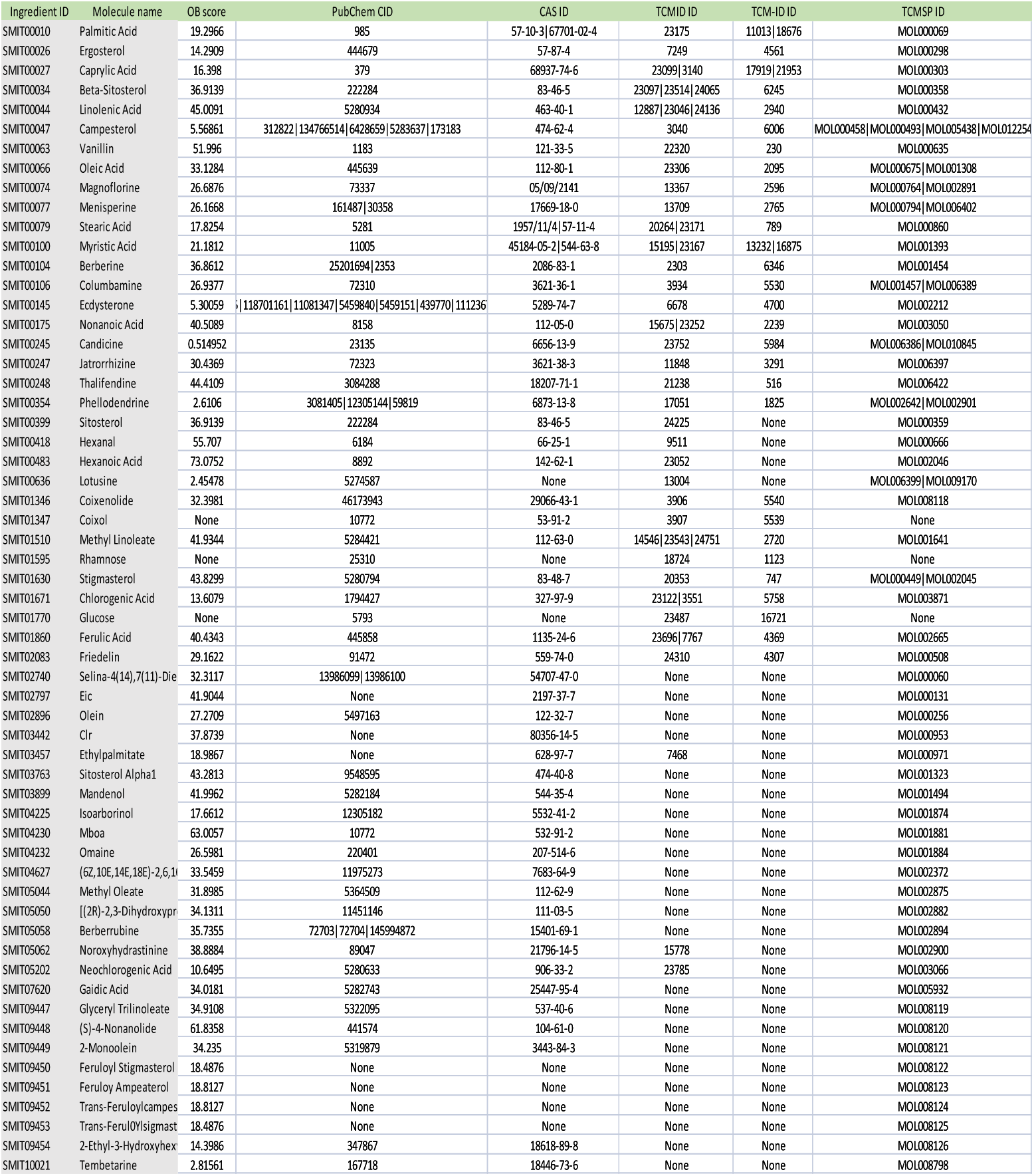
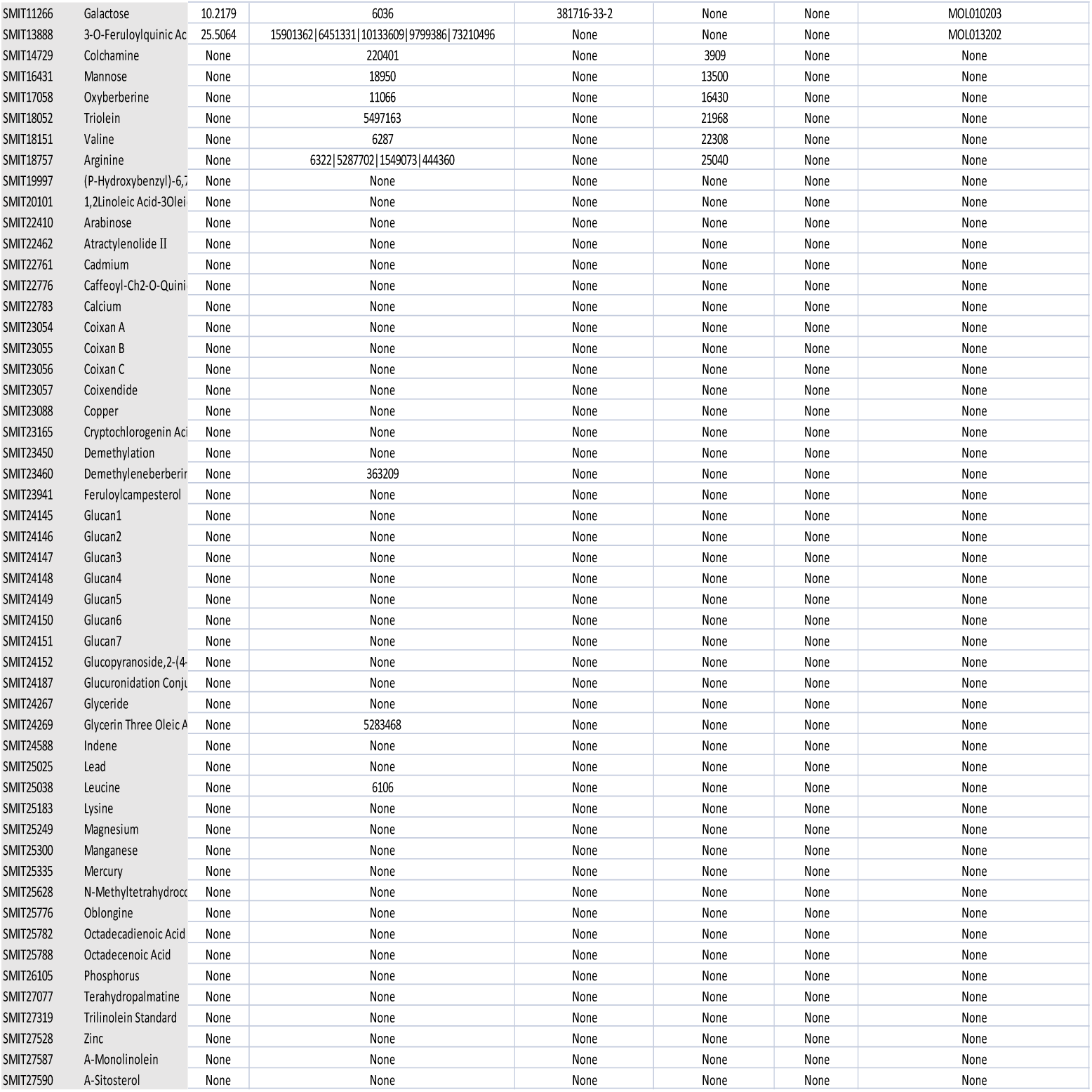

**Supplementary Table S2.**
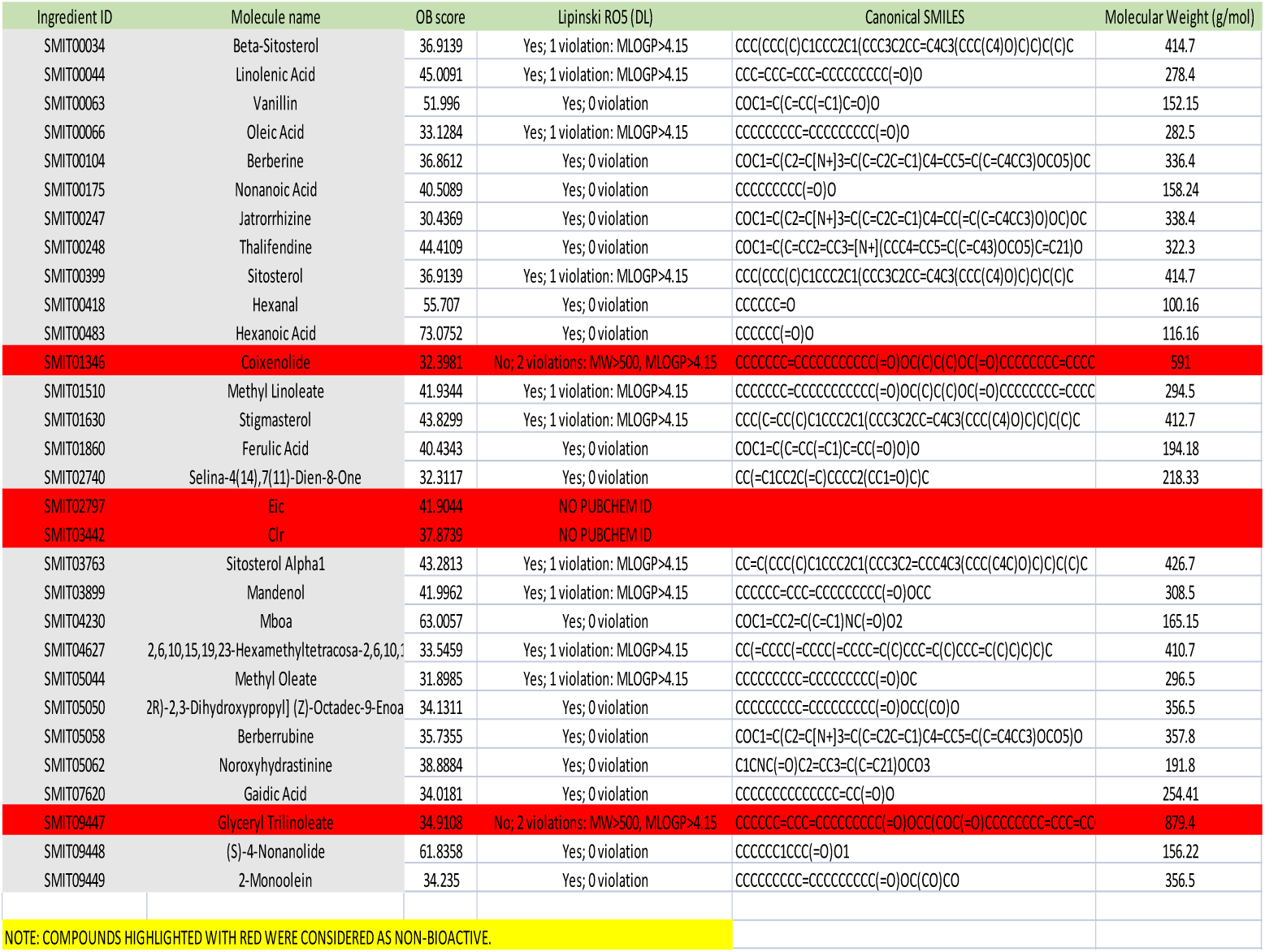

